# What is the evidence available to support our knowledge about threats to the conservation of Galliformes in the Greater Himalaya?

**DOI:** 10.1101/2021.02.23.432604

**Authors:** Garima Gupta, Matthew Grainger, Jonathon C. Dunn, Roy Sanderson, Philip J.K. McGowan

**Affiliations:** School of Natural and Environmental Sciences, Agriculture Building, King’s Road, Newcastle upon Tyne NE1 7RU; Norwegian Institute for Nature Research, Trondheim, Sør-Trøndelag, Norway; Institute of Neuroscience, Henry Wellcome Building, The Medical School, Framlington Place, Newcastle University, Newcastle upon Tyne, NE2 4HH

**Keywords:** Conservation, Extinction, Galliformes, Himalaya, Literature Review, Pheasants, Threats

## Abstract

Biodiversity is at a heightened risk of extinction and we are losing species faster than any other time. It is important to understand the threats that drive a species towards extinction in order to address those drivers. In this paper, we assess our knowledge of the threats faced by 24 Himalayan Galliformes species by undertaking a review to identify threats reported in the published literature and the supporting evidence that the threat is having an impact on the species population. Only 24 papers were deemed suitable to be included in the study. We found that biological resource use, agriculture and aquaculture are the predominant threats to the Galliformes in the Greater Himalaya but the evidence available in the studies is quite poor as only one paper quantified the impact on species. This study shows that major gaps exist in our understanding of threats to species and it is imperative to fill those gaps if we want to prevent species from going extinct.

## 2. Introduction

There is increased political realisation of the societal impacts of deteriorating biodiversity (Griggs *et al*., 2013; IPBES 2019). This is encapsulated in a variety of multilateral environmental agreements (MEAs), most notably the Convention of Biological Diversity (CBD), and in the UN Sustainable Development Goals, and national policies and strategies. The two main factors behind species extinction are continual growth in both human population and increase in per capita consumption (Pimm *et al*., 2014; Guerry *et al*., 2015). These give rise to a variety of pressures that have direct consequences for species and the scale of these pressures is increasingly understood.

General patterns in the intensity and distribution of these pressures can be drawn from the IUCN Red List of Threatened Species. One of the most significant anthropogenic pressures is agricultural activity, with 62% (5407) of those species that have been assessed as threatened or near threatened affected by crop farming, livestock farming, timber plantation, and/or aquaculture (Maxwell *et al*., 2016). Overexploitation of species for consumption by humans has been long considered to be a significant threat to many species (Fa *et al*., 2003; Milner-Gulland & Bennett, 2003; Vié *et al*., 2009; Wittemyer *et al*., 2014). Some species may also be overexploited for non-subsistence purposes, such as trade or recreation and there are many high profiles cases, for example tigers (*Panthera tigris*), which are classified as Endangered, and are hunted illegally because of the high commercial demand for its skin and bones. Often species are threatened by multiple threats, with the combined effects of overexploitation and agricultural activity having the greatest impacts on biodiversity (Mace *et al*., 2000; Peres, 2001). Together they have contributed to 75% of extinctions since AD 1500 (Maxwell *et al*., 2016).

Pressures on biodiversity may increase or decrease over time, and this may be over the short or long-term, and new pressures may emerge. As pressures change, the specific threats that they produce and negative impacts that they have on species, and indeed other elements of biodiversity, will also change. Therefore, to identify the most appropriate conservation measures in a given place and time, whether policy, legislation, management, or some other intervention, we do need to know that the conservation action will have a beneficial impact on species.

Aichi Biodiversity Target 12 stated that ‘by 2020, the extinction of known threatened species has been prevented and their conservation status, particularly of those most in decline, has been improved and sustained’ (CBD, 2010). To halt extinctions and improve the conservation status of species we need to go beyond simply understanding species extinction risks, and knowledge of pressures and their scale, and move towards detailed understanding of how to mitigate threats so that species can recover. In other words, we need to deepen our assessments of pressure and the conservation status of species so that we know which threats have a documented impact on species’ populations and where, so that, when they are reduced, they can result in population increases. In this paper, we explore what we know about threats to a group of 24 bird species, the Galliformes of the Himalaya.

Galliformes are important ecologically, economically, and culturally in the Himalaya and are one of the most threatened bird orders (McGowan & Fuller, 2006; Sathyakumar & Sivakumar, 2007) and yet, no study specifically examines all threats facing an entire taxonomic group within the Himalaya. Most studies to date have focussed on only a few species, and we need to be clear about the impact of a reported threat on the population of a species. To make optimal use of limited conservation resources, we need to know with as much certainty as possible what the threats are, where they occur, and whether there are any patterns in the type and spatio-temporal distribution of threats for Himalayan Galliformes. This should then form the basis of targeted responses.

There is a need to understand what is really known, rather than assumed, about the impacts of threats on species for which there is little extant information on their ecology, behaviour, or life-history. Where there is no firm information on how threats are affecting species and what is needed to address these threats, we need to structure our predictions logically and transparently (e.g. Grainger *et al*. (2018)). An objective approach must be taken to increase our understanding of threats to Galliformes where the quality of published evidence that a threat results in population decline is variable. One approach is to review the habitats of species that have been assessed (i.e. their IUCN Red List status) and determine whether there are patterns in changes to these habitats, and their associated species, which could guide conservation interventions to maximise their benefits. This would help make conservation responses logical and transparent, as it would be clear when the impacts of threats are being inferred from other information.

In this paper, we seek to understand our knowledge of the threats facing Himalayan Galliformes. We do this by undertaking a literature search to identify the threats reported in the literature and the evidence supporting them.

## 3. Methods

### 3.1 Assessing published knowledge of threats to Himalayan Galliformes

#### 3.1.1 Search engine and search terms

Searches were undertaken on the Web of Science core collection and Google Scholar for research articles that included potential threats to Galliformes in the Himalaya. Search terms were selected to increase the possibility of obtaining relevant articles on all potential threats. The main aim of the literature search was to glean information on possible factors thought to cause declines in Galliformes in the Himalayan region, and what empirical evidence existed for these factors actually causing declines in species’ populations. The term “Galliformes” tends to be used in keywords of papers, if not in the paper themselves, to describe the taxonomic group to which each species belongs.

Web of Science was searched for terms “TS = ((galliform* OR pheasant OR partridge OR quail) AND threat*)”, “TS =((galliform* OR pheasant OR partridge OR quail) AND Himalaya*)” and “TS =((galliform* OR pheasant OR partridge OR quail) AND Himalaya* AND threat*)” and “TS =((galliform* OR pheasant OR partridge OR quail) AND Himalaya* AND conserv*)” and “TS =((galliform* OR pheasant OR partridge OR quail) AND conserv*)”. Google Scholar was also searched for “threats to Galliformes in the Himalaya”. Articles from “Proceedings of the 3^rd^ International Galliformes Symposium, 2004” (Fuller and Browne, 2005), which was a CD-ROM and so the articles not easily indexed, were also screened.

Papers from Environmental Sciences/Ecology fields were searched for inclusion in the study since there was an overlap of research articles in other fields. These fields have been identified in the Web of Science database, but Google Scholar does not provide these fields to narrow down the search results. Searches were made across all years and the language search criterion was set to include papers in English.

#### 3.1.2 Criteria for inclusion in study

All papers were screened based on titles and abstracts. The primary inclusion criteria were a) studies should only focus on Himalayan Galliformes species; b) papers should only be primary literature i.e. no reviews, unpublished reports or action plans; c) studies should be within the Himalayan countries of India, Pakistan, China, Nepal and Bhutan. Articles that dealt with other species and were outside the Himalayan region were discarded.

### 3.2 Quality of threat reporting, and definitions used in classification of quality of documentation of threats

Papers included in the study were assigned to one of four categories according to the evidence that the paper provided for each threat that it reported. The four categories were:

#### a) Unsubstantiated Assertion

A study was categorised as ‘unsubstantiated assertion’ when a threat was reported as a probable factor in driving a species towards population decline but the threat had not been documented in the study site.

#### b) Threat Documented

A study was allocated to this category when a threat had been documented but there was no evidence to show that the threat was causing a decline in species’ numbers.

#### c) Impact Inferred

A paper was categorised as ‘impact inferred’ if it showed that a threat did exist and then suggested that the threat has had an impact on a Galliformes species but did not provide evidence to show what that impact was in the paper.

#### d) Impact Documented

A study was classified as ‘impact documented’ when there was direct evidence to show that the population had declined due to a reported threat.

To avoid any biases in categorising papers, two independent authors reviewed all papers separately and classified them to one of the four categories. 24 papers were reviewed by three authors. We assessed the proportion of agreement by calculating Cohen’s Kappa with Psych package (Revelle 2019) in R version 3.6.1.

### 3.3 Threats reported to Himalayan Galliformes in published literature and their classification

Threats reported in research papers included in the study were identified and then classified based on Level 1 categories of the International Union for Conservation of Nature-Conservation Measures Partnerships unified Classification of Direct Threats (IUCN CMP, 2019) (see Supplementary Table 1). The Level 1 categories in the IUCN threat classification are: Biological Resource Use, Agriculture and Aquaculture, Natural System Modifications, Residential, Transportation and Service Corridors, Human Intrusion and Disturbance, Pollution and Others. The papers found during the literature survey were nearly all published before the Classification of Direct Threats was adopted and so they did not report threats using the terminology of the Level 1 categories of IUCN threat classification. The way that the papers reported each threat to a species made it straightforward to classify the threats in one of the Level 1 categories.

## 4. Results

### 4.1 Assessing published knowledge of threats to Himalayan Galliformes

The total number of papers identified by searching the Web of Science for “TS = ((galliform* OR pheasant OR partridge OR quail) AND threat*)” were 181 results. Similarly “TS = ((galliform* OR pheasant OR partridge OR quail) AND Himalaya*)” and “TS = ((galliform* OR pheasant OR partridge OR quail) AND Himalaya* AND threat*)” and “TS = ((galliform* OR pheasant OR partridge OR quail) AND Himalaya* AND conserv*)” and “TS = ((galliform* OR pheasant OR partridge OR quail) AND conserv*)” returned 36, nine, 22, and 620 results respectively. Google Scholar returned 667 results when the term “threats to Galliformes in the Himalaya” was used. Duplicate papers that were returned from different database searches were eliminated. Another two papers were included from “Proceedings of the 3rd International Galliformes Symposium 2004” (Fuller and Browne, 2005). (See Figure 1 for details).

**Figure 1:**
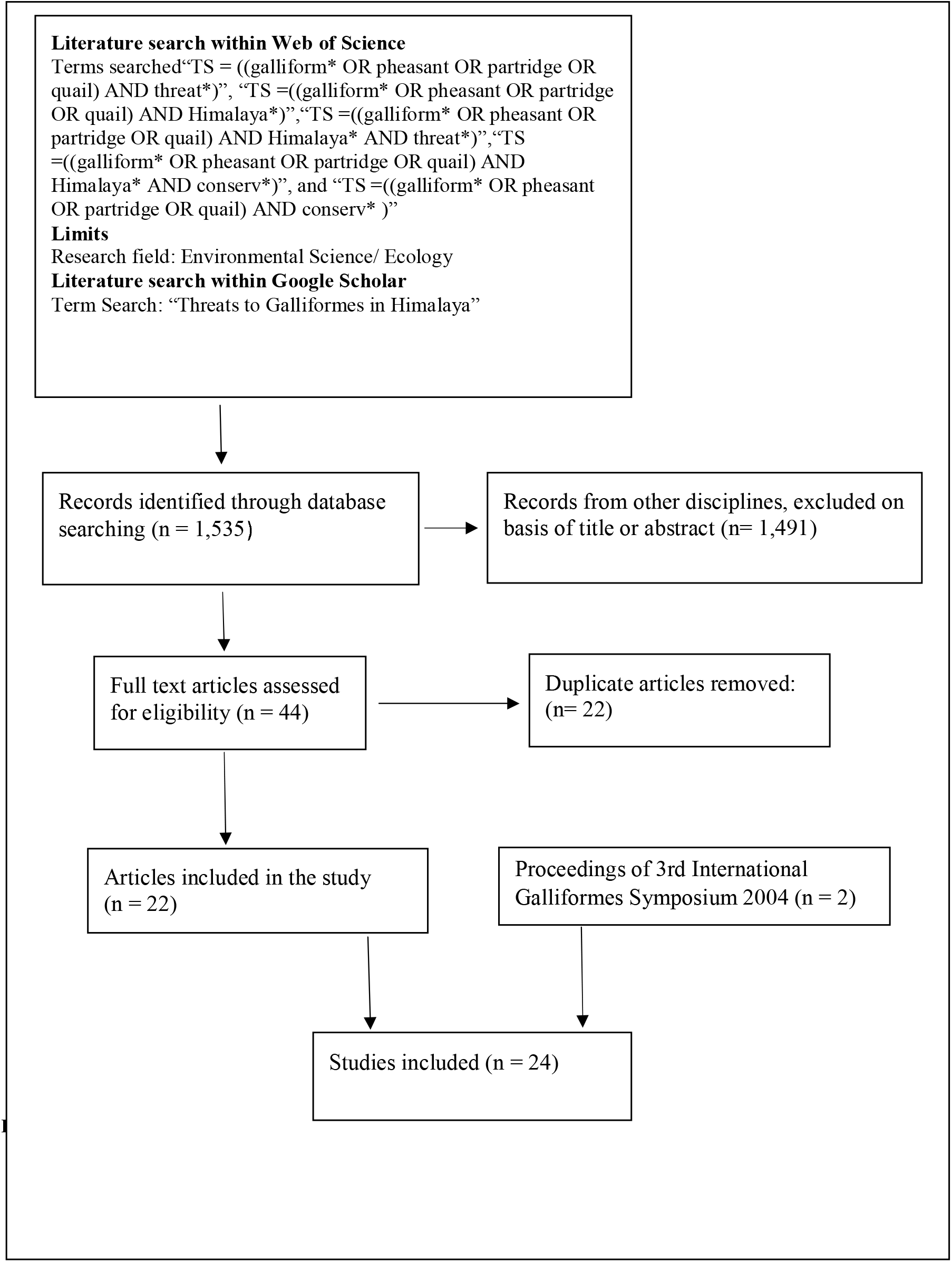
PRISMA flow diagram of literature search, based on Liberati *et al*. (2009).

The searches returned a total of 1,535 unique references of which only 22 (1.4%) met the inclusion criteria and were consequently included in the study. Approximately 97% (1,491) references were excluded as they did not fit the inclusion criteria and were, for example based on lab-based genetic and molecular studies, which has no relevance to the current study.

### 4.2 Quality of threat reported

Papers were assessed for the quality of threat reporting and of the 24 studies identified, only one paper quantified the effect of hunting on the population of the Himalayan Galliformes (see Figure 2). Sixteen papers (64%) included in the study reported threats based on unsubstantiated assertion i.e. that is there is no substantive evidence to prove that the threat documented was found in the study area. The number of papers classified under threat documented and impact inferred are four and three respectively. There was a high agreement between all reviewers in classifying the papers (Cohen’s Kappa = 0.83, 95% CI 0.63-0.83).

**Figure 2.**
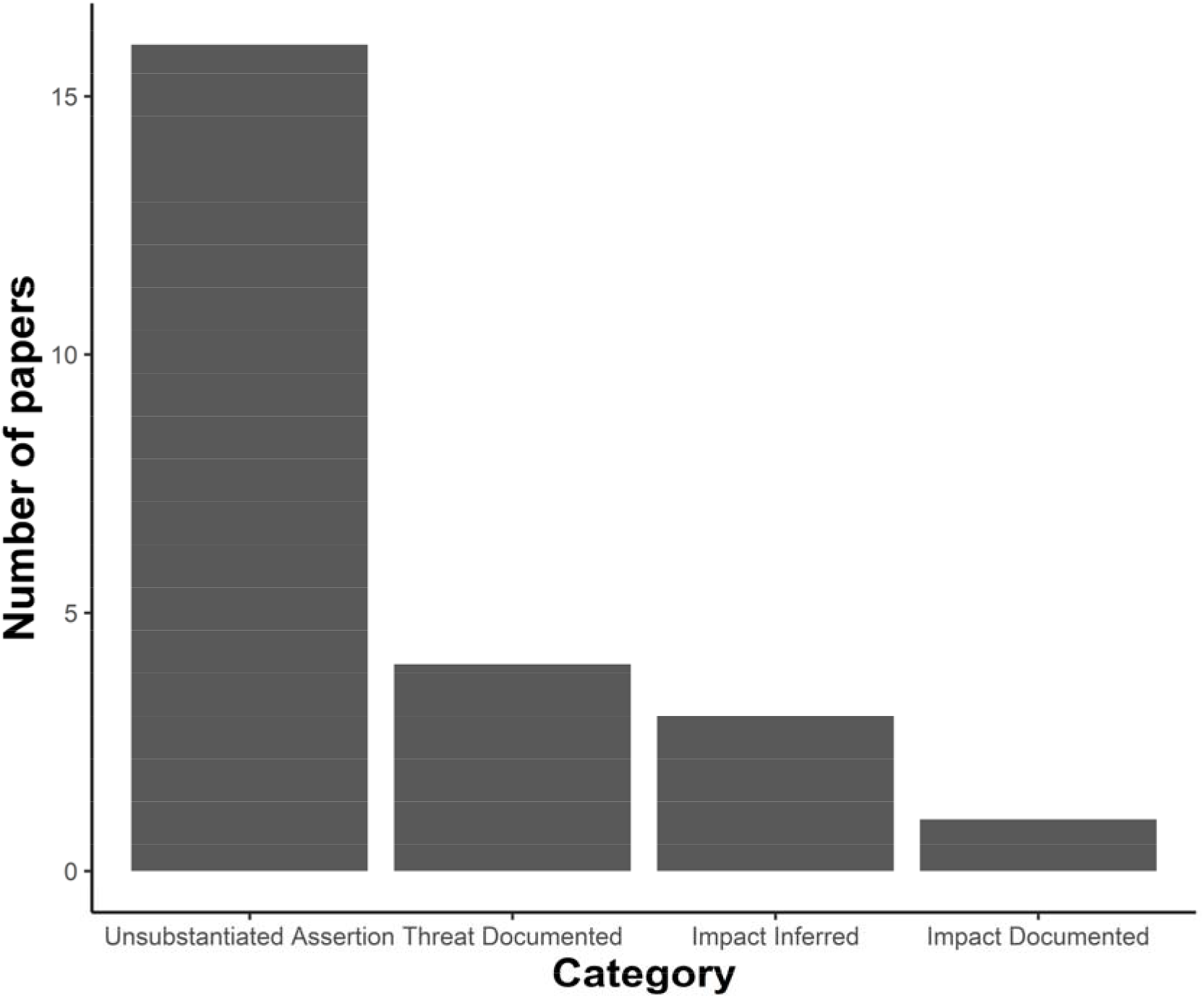
Nature of the evidence reporting threats to 24 Galliformes in the Greater Himalaya in 24 studies in the peer reviewed literature.

### 4.3 Threats reported to the Himalayan Galliformes

Some papers reported more than one threat to the Galliformes in the Greater Himalaya, which meant that there were 35 reported threats in the 24 papers (see Figure 3).

**Figure 3.**
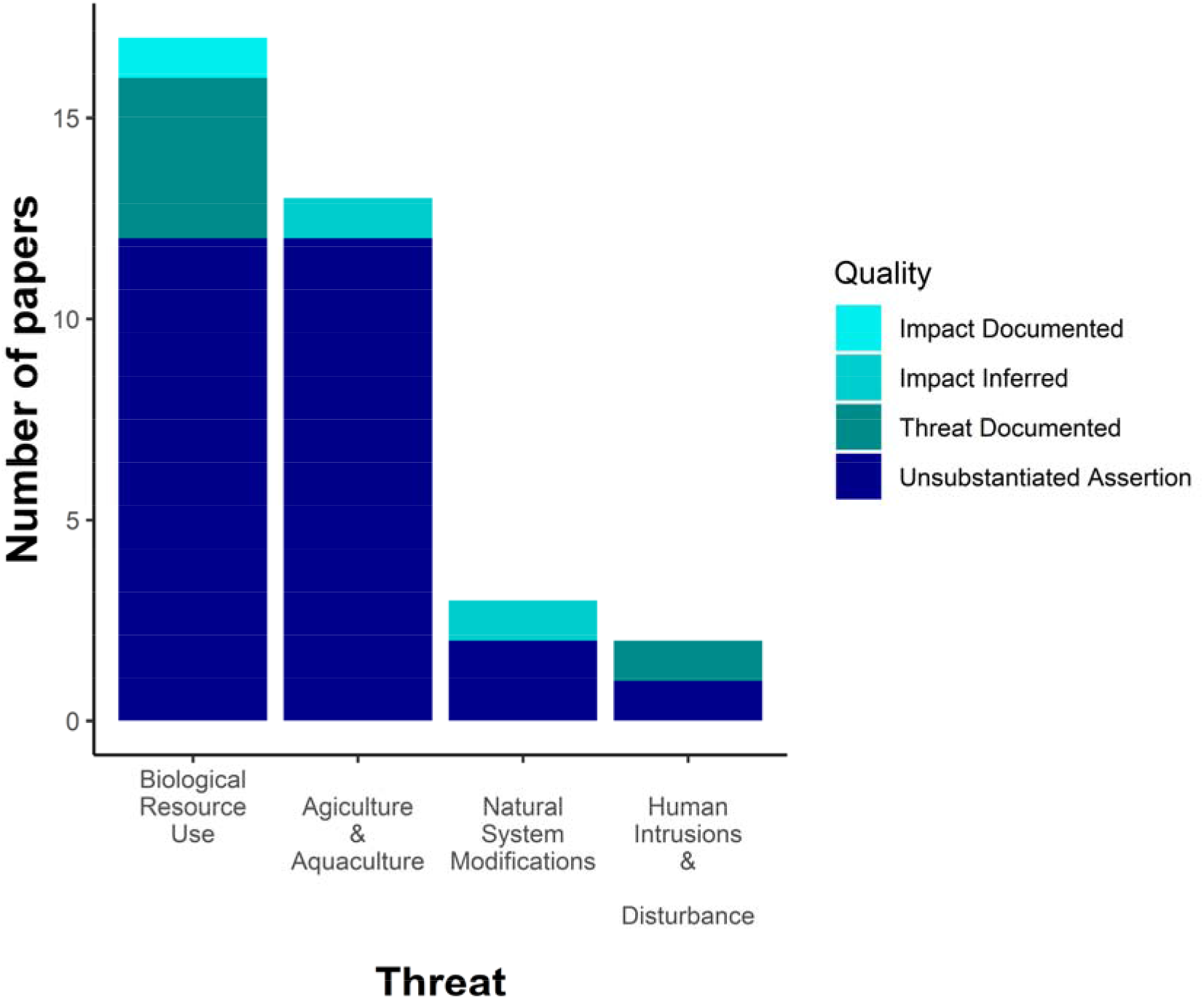
Different types of threats reported in research papers included in the study and the quality of documentation of threats.

Sixteen papers reported Biological Resource Use as a potential threat to Himalayan Galliformes (see Figure 3). Of these 16 papers, only one paper documented Biological Resource Use as an impact, whilst the majority of them were unsubstantiated assertions. Agriculture and Aquaculture was reported in 13 papers, out of which one was classified under threat inferred and rest all were unsubstantiated assertions. Development activities such as hydroelectric dams categorised under Natural System Modification were also reported as a threat to the Himalayan Galliformes.

## 5. Discussion

Effective conservation decision-making is challenging because our knowledge of the natural world is imperfect and the impact of our actions upon it are uncertain (Bolam *et al*., 2018). It is not easy to predict the impact of conservation actions on each species, and it is also a challenge to determine where and how to act to ensure maximum long-term conservation benefits (e.g. Grainger *et al*. (2018)). In this study, ‘only’ 24 papers from a total of 1,535 Galliformes studies were found that reported threats to the Galliformes of the Greater Himalayan region. Sixteen papers had a threat reported but provided no firm evidence that it was operating in the area studied and only one paper had firm, documented evidence that a threat was having an impact on a population. Biological Resource Use and Agriculture & Aquaculture were reported as main pressures on Himalayan Galliformes.

Despite being a highly threatened group of birds with 25% of the 308 Galliformes species threatened with extinction (McGowan, 2002; Grainger *et al*., 2018), the group remains understudied. This incomplete knowledge is reflected by only 24 papers documenting impacts of threats on a Galliformes species that are causing population declines. This suggests that there is a need for both field studies in the region to study human pressures on the species, and a change in the way studies examine and report threats and their impact on species.

Hunting and poaching, which is classified under Biological Resource Use (see Supplementary Table 1), was found to be the predominant threat reported with 16 papers reporting hunting as a threat to Galliformes in the Greater Himalaya. Even though hunting and poaching is prohibited in many countries, many species are still hunted for their body parts and meat. Many tropical areas suffer from hunting that can have profound impacts on biodiversity, which can then have negative cascading effects on wider food webs and ecosystems (Milner-Gulland & Bennett, 2003; Bennett *et al*., 2007; Wright *et al*., 2007). Although, wildlife in Asia has been undergoing rapid declines in geographic range and population, there are relatively few studies that have documented the actual impact of hunting as a problem for a species (O’Brien *et al*., 2003; Steinmetz *et al*., 2006; Corlett, 2007). Thus, there is often not enough evidence to determine the significance of hunting in the decline of individual species. Of the 16 papers that reported hunting as a threat to the Galliformes in the Greater Himalaya, only one had the threat properly documented, whereas others made unsubstantiated assertions i.e. there was no evidence to prove that hunting was a threat to the Galliformes.

Galliformes are hunted for food to varying degrees throughout their geographical range. Hunting of animals is illegal in many countries and this might be one of the reasons behind lack of evidence on hunting in the Himalayan area. People might not be open about the prevalence of hunting of the animals in the region, as they might be afraid of being caught and penalised for their actions. Other threats include habitat loss due to deforestation activities mainly for agriculture such as *jhum* cultivation (slash and burn). Thirteen papers reported Agriculture and Aquaculture, which includes threats from farming and ranching as a result of agricultural expansion (see Supplementary Table 1) as the second biggest threat to the Galliformes. Since the Greater Himalaya has the most extensive areas of glaciers and permafrost globally and is the source of nine large rivers, it is called ‘the water tower of Asia’ (Xu et al., 2009; Xu & Grumbine, 2014). This makes the Himalaya a potential source for production of hydroelectric energy resulting in deforestation and submergence of a huge area, with subsequent loss of species habitat.

There is therefore a need to understand threats to biodiversity, identify regions where risks occur and to quantify the rates of change in those threats, in order to ensure that conservation actions are appropriately targeted and be most effective to achieve long-term environmental goals (Geldmann *et al*., 2015). We can achieve this by focussing research on threats in areas with high biodiversity and high human pressures whilst ensuring that the research is designed and reported to a high standard. In conclusion, this study has identified major gaps exists in our knowledge on the threats to species that can lead to extinction. It is imperative to fill these gaps if we are to achieve the successor to the Convention on Biological Diversity’s Aichi Target 12 of halting species extinction and improving the status of the declining threatened species, which is being negotiated at present.

## Recommendations

- The way a threat is reported in any study needs to be supported by empirical evidence. If a threat has been identified in the area and if the documented threat results in decline of a species population then only the direct negative results of that treat should be reported, whilst other reported impacts should be treated with caution.
- We need to design studies to directly assess threats rather than infer them from circumstantial evidence. This will be difficult, but there is a pressing need to design better observational studies (and pseudo-experimental designs), and better social-ecological studies to assess this directly. Studies on population parameters are needed, for example survival could be monitored through telemetry. We should also make use of integrated population models that use data on populations, survival and reproduction and combine these to reconstruct population dynamics - these simulations can then lead to inference about the influence of poaching on population persistence over time.
- Studies on specific species could be coordinated so that key components of the population parameters are assessed by different researchers and then combined into a single integrated population model. For example, IUCN Species Survival Commission Galliformes Specialist Group (https://www.iucn.org/commissions/ssc-groups/birds/galliforme) could coordinate this for the Himalayan Galliformes.

## Author contributions

Study design and data collection: GG, MG, JCD, RS, and PJKM; data analysis: GG, MG, PJKM; and writing and revision: all authors.

## Acknowledgements

This research received no specific grant from any funding agency, or commercial or not-for-profit sectors.

## Conflicts of interest

None.

## Ethical standards

This research abided by the Bird Conservation International guidelines on ethical standards.

## Notes

### Competing Interest Statement

The authors have declared no competing interest.

