## Supplementary Table 1 for "What is the evidence available to support our knowledge about threats to the conservation of Galliformes in the Greater Himalaya?"

**Supplementary Table 1. IUCN-CMP unified level 1 classification of direct threats (IUCN CMP 2019).**

| **Level 1 classification of threats** | **Examples** |
| --- | --- |
| **Residential & Commercial Development** | Threats from human settlements or other non- agricultural land uses with a substantial footprint |
| **Agriculture & Aquaculture** | Threats from farming and ranching as a result of agricultural expansion and intensification, including silviculture, mariculture and aquaculture (includes the impacts of any fencing around farmed areas) |
| **Energy Production & Mining** | Threats from production of non-biological resources |
| **Transportation & Service Corridors** | Threats from long narrow transport corridors and the vehicles that use them include associated wildlife mortality |
| **Biological Resource Use** | Threats from consumptive use of "wild" biological resources including both deliberate and unintentional harvesting effects; also persecution or control of specific species |
| **Human Intrusions & Disturbance** | Threats from human activities that alter, destroy and disturb habitats and species associated with non-consumptive uses of biological resources. |
| **Natural System Modifications** | Threats from actions that convert or degrade habitat in service of “managing” natural or semi-natural systems, often to improve human welfare |
| **Invasive & Other Problematic Species, Genes & Diseases** | Threats from non-native and native plants, animals, pathogens/microbes, or genetic materials that have or are predicted to have harmful effects on biodiversity following their introduction, spread and/or increase in abundance |
| **Pollution** | Threats from introduction of exotic and/or excess materials or energy from point and nonpoint sources |
| **Geological Events** | Threats from catastrophic geological events |
| **Climate Change & Severe Weather** | Threats from long-term climatic changes which may be linked to global warming and other severe climatic/weather events that are outside of the natural range of variation, or potentially can wipe out a vulnerable species or habitat |
| **Other Options** | The threats classification scheme is intended to be comprehensive, but as there are often new and emerging threats, this option allows for these new threats to be recorded |
